# Anti-NMDAR encephalitis alters intrinsic spatiotemporal coding by enhancing neuronal coupling and clustering

**DOI:** 10.1101/2024.08.20.608793

**Authors:** Vahid Rahmati, Jürgen Graf, Mihai Ceanga, Dario Cuevas Rivera, Holger Haselmann, Sabine Liebscher, Harald Prüss, Knut Holthoff, Knut Kirmse, Christian Geis

**Author notes:** Correspondence **Address for correspondence:** Christian Geis, MD Department for Neurology Section Translational Neuroimmunology Jena University Hospital Am Klinikum 1, D-07747 Jena, Germany. These authors contributed equally (Vahid Rahmati, Jürgen Graf, Mihai Ceanga).

## Abstract

Autoimmune anti-NMDA-receptor encephalitis is characterized by severe neuropsychiatric symptoms including memory dysfunction and seizures. However, it remains enigmatic what functional changes at the multi-neuronal level mediate network dysfunction. We used two-photon *in vivo* recording in a passive-transfer mouse model with patient’s monoclonal anti-GluN1-autoantibodies during slow-wave sleep-like conditions, a critical phase for memory processing. We find enhanced functional coupling and clustering between hippocampal CA1 pyramidal neurons (PNs), which intrinsically expose the network to hypersynchrony. These connectivity changes are associated with a selective preservation of strong excitatory synapses despite overall reduced excitation, thus enhancing hub-like properties of functionally connected PNs. Furthermore, we find abnormal PN firing characteristics, decreased transmission failure, and increased similarity of spontaneous spatiotemporal activity patterns, all affecting CA1 intrinsic neuronal coding. Collectively, the functional rewiring of hippocampal networks and altered intrinsic information processing provide new mechanistic insights into the NMDAR-hypofunction consequences and pathomechanisms of anti-NMDAR encephalitis symptomatology.

## Introduction

NMDA receptors (NMDARs) play a pivotal role in excitatory transmission and both long-term synaptic depression (LTD) and potentiation (LTP), subserving memory, learning, and psychosocial behavior (Lau and Zukin, 2007; Paoletti et al., 2013; Papi et al., 2024). Hypofunction of NMDARs is linked to various neuropsychiatric disorders, such as schizophrenia and Alzheimer’s disease (Lau and Zukin, 2007; Paoletti et al., 2013; Balu, 2016; Liu et al., 2019). Anti-NMDAR encephalitis is a severe autoimmune disorder, caused by autoantibodies targeting the NR1 (GluN1-Ab) subunit of the NMDAR (Dalmau et al., 2017). The affected patients share phenotypical similarities to schizophrenia and suffer from severe neuropsychiatric symptoms, ranging from psychosis and cognitive deficits to severe disruption of sleep/wake cycles, dysautonomia, and seizures (Dalmau et al., 2017; Papi et al., 2024). The current standard immunotherapy is often insufficient in managing the symptoms in severe cases (Sell et al., 2021; Papi et al., 2024). GluN1-Ab predominantly reduces neuronal surface expression of hippocampal NMDARs, thus disrupting NMDAR currents and LTP (Hughes et al., 2010; Planagumà et al., 2016; Ladépêche et al., 2018). Despite the impaired synaptic excitatory signaling (Wright et al., 2021; Ceanga et al., 2023; Steinke et al., 2023), favoring inhibition dominance, cortico-hippocampal circuits under GluN1-Ab are prone to hypersynchronous discharges whose underlying mechanisms remain enigmatic.

By promoting synaptic plasticity and facilitating cortical–hippocampal dialogue, slow-wave sleep (SWS, also called non-REM sleep) plays a pivotal role in learning and memory processing (Walker, 2009; Mitra et al., 2016; Tang et al., 2017; Tamaki et al., 2020), which are often impaired in anti-NMDAR encephalitis (Dalmau et al., 2017; Papi et al., 2024). During SWS, which accounts for most of the sleep time, hippocampal networks exhibit a rich repertoire of internally generated activity patterns, which largely recapitulate sequences of neuronal activation for places or events experienced/learned in the awake state (‘neural replays’) to stabilize memories (Wilson and McNaughton, 1994; Foster, 2017; Lewis et al., 2018). These patterns are mainly based on functional neuronal coupling, mediating co-activated or sequentially activated neuronal ensembles (or clusters), which play an important role in processes such as memory, perception and behavior (Wilson and McNaughton, 1994; Buzsáki, 2004; Luczak et al., 2007; Alejandre-García et al., 2022; Yuste et al., 2024). Furthermore, their alterations were proposed to underlie brain dysfunction in neurological diseases, including schizophrenia (Northoff and Duncan, 2016; Hamm et al., 2017; Sauer and Bartos, 2022). We hypothesize that impairments in these internal building-blocks of neural computations underlie learning and memory dysfunction in condition of NMDAR hypofunction, as in anti-NMDAR encephalitis.

Using *in vivo* two-photon Ca^2+^ imaging in the passive-transfer anti-NMDAR encephalitis mouse model (Planagumà et al., 2016) under anesthesia, thereby minimizing sensory input through emulating SWS-like sleep in the CA1 hippocampal microcircuit, we sought to address: I) What change in functional circuitry at multi-neuronal level mediates network hypersynchrony despite disrupted synaptic excitation? II) How does this alteration affect intrinsic spatiotemporal coding possibly linked to impaired learning and memory phenotypes of the disease? We find that GluN1-Ab suppresses overall network activity but concomitantly drives it hypersynchronous through enhanced functional coupling and clustering between pyramidal neurons (PNs). We further find an abnormal alteration of spatiotemporal patterns and neuron-to-neuron communication. Overall, the functional rewiring of hippocampal networks and altered intrinsic information processing provide new mechanistic insights into the NMDAR-hypofunction consequences and pathomechanisms of anti-NMDAR encephalitis.

## Results

We employed the established passive-transfer mouse model with *in vivo* chronic intraventricular delivery of pathogenic immunoglobulin G (IgG) via osmotic pumps (Planagumà et al., 2016), applying either NMDAR encephalitis patient-derived monoclonal NMDAR-GluN1 subunit (GluN1-Ab) or control IgG (Ctrl-Ab), for 14-16 days (**Fig. 1A**, left). We applied high-speed acousto-optic two-photon imaging to record somatic Ca^2+^ transients (CaTs), from individual CA1 neurons in *stratum pyramidale*, mainly comprising glutamatergic PNs (**Fig. 1A**, right) (Graf et al., 2022). Experiments were conducted in head-fixed mice under isoflurane anesthesia (∼0.6%) to induce SWS-like sleep that in many ways resembles non-REM natural sleep (Luczak et al., 2007) and minimizes sensory input to the cortico-hippocampal circuitry. For each mouse, one field of view (FOV) was recorded for about one hour, subjected afterwards to an offline cell-detection procedure (**Fig. 1B**) to extract fluorescence traces per PN (**Fig. 1C**), followed by the reconstruction of CaT onsets, as a proxy for firing activity (**Fig. 1D** top). In addition, we conducted LTD recordings from CA1 region *ex vivo*. Full statistical results are provided in **Table S1**.

**Figure 1.**
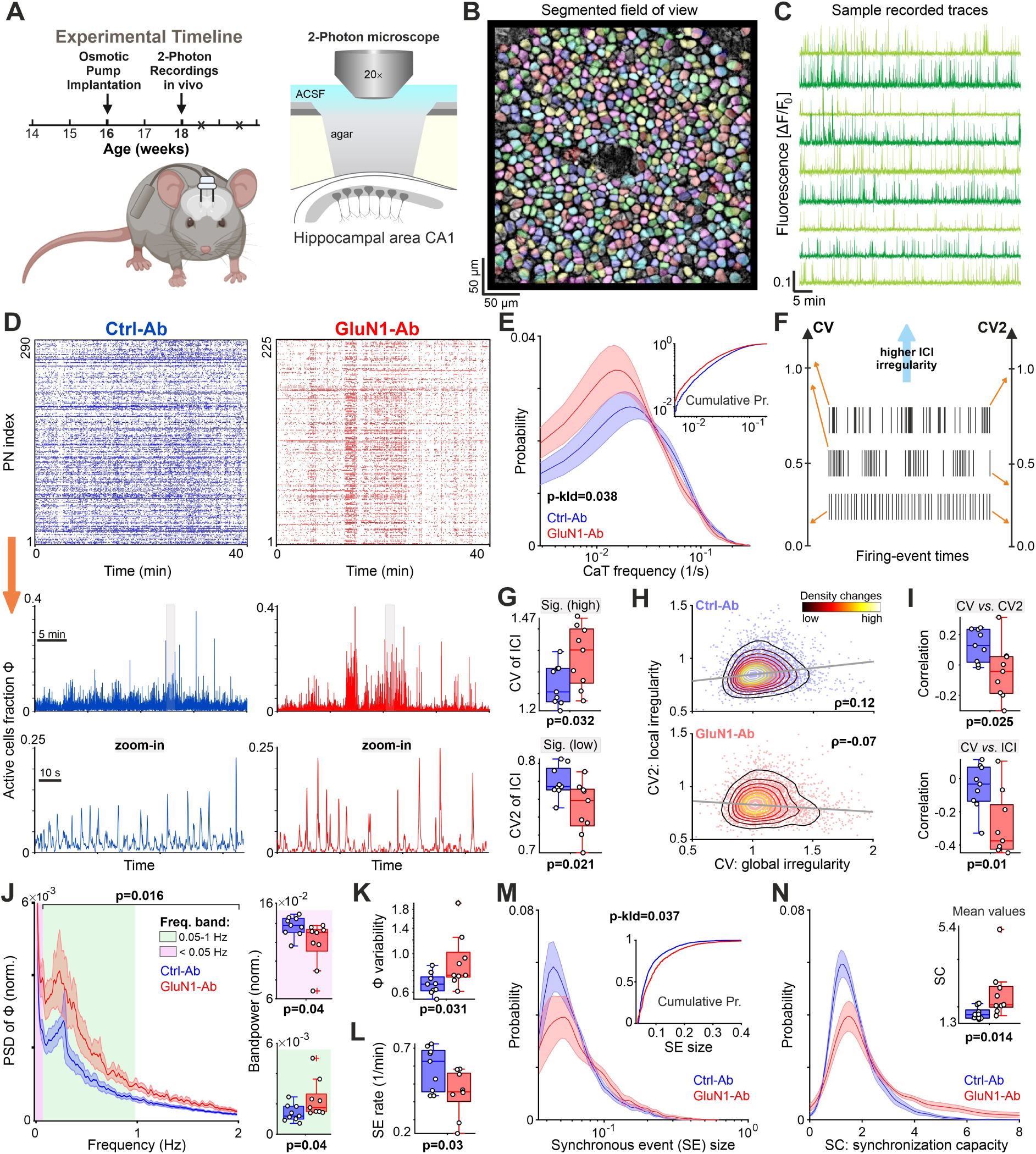
GluN1-Ab augments CA1 network synchrony while reducing overall activity. (**A**) Schematic of the experimental timeline (left) and recording configurations (right). (**B**) Representative field of view (FOV) of 2-Photon recordings from CA1 area using OGB1-indicator *in vivo*, overlaid by the detected region of interests (ROIs) representing the somata of putative pyramidal neurons (PNs). (**C**) Timeseries of Ca^2+^ fluctuations extracted from example ROIs, here PNs. (**D**) Top: rasterograms of reconstructed onsets of PNs’ Ca^2+^ transients (CaT, dots) for two example FOVs (1 FOV per mouse). Middle: the corresponding timeseries of active PNs fraction Φ(t). Bottom: the zoom-in of periods marked in middle row (gray rectangle). (**E**) Distributions of mean CaT frequency of PNs (n=9 mice/4729 cells in Ctrl-Ab; n=9 mice/3200 cells in GluN1-Ab). The p-kld denotes the p-value of permutation test based on the Kullback-Leibler divergence (KLD) used to compare the overall shape of distributions. Inset: cumulative probability of CaT frequencies of all mice (concatenated per group). (**F**) Schematic of global (CV) and local irregularity (CV2) measures of inter-CaT intervals (ICIs). (**G**) Irregularity of PNs with significantly high CV or low CV2 values. (**H**) Negative relationship between CV and CV2 of ICIs under GluN1-Ab. ρ indicates the average Spearman’s rank correlation coefficient over those computed per mouse. Each dot represents a PN. PNs of all mice are shown (Ctrl-Ab: 4729, and GluN1-Ab: 3200 cells). The contour lines indicate the slope of cell-density changes. (**I**) The relationship between global and local irregularity (top) and median-ICI of PNs (bottom). Spearman’s rank correlation coefficient was computed separately per FOV (i.e. mouse). (**J**) Power spectral density of Φ(t) (PSD, left), and corresponding bandpowers. PSD was normalized to total power per mouse. (**K**) Variability of network activity fluctuations computed as coefficient of variation of Φ(t). (**L**) The occurrence rate of synchronous events (SEs). (**M**) Redistribution of SE size towards larger values under GluN1-Ab. Same format as in (E). (**N**) Increased synchronization capacity of PNs, i.e. their tendency to participate in SEs, under GluN1-Ab. Curves presented as mean ± SEM. For statistical information, see Table S1.

### GluN1-Ab alters firing rates and patterns of CA1 PNs

First, we investigated GluN1-Ab effects on the firing characteristics (frequency and structure) of CA1-PNs (**Fig. 1D-I**). This analysis revealed an overall alteration in the CaT frequency distribution, with an increase in neurons with lower activity rather than a mean shift across the entire spectrum (**Fig. 1E, Table S1**). GluN1-Ab tended to increase the global irregularity (CV) but to diminish temporally local irregularity (CV2) of inter-CaT intervals (ICIs; **Figs. 1F-I, S1A-C**). Specifically, compared to surrogate data obtained by randomizing CaT events per neuron, PNs with a significantly high CV showed even higher CV levels, while PNs with a significantly low CV2 exhibited more decreased CV2 levels under GluN1-Ab (**Fig. 1G**). This was further reflected in the correlation between CV and CV2 of ICIs, which reduced from positive under Ctrl-Ab to negative values under GluN1-Ab (**Fig. 1H,I**). These observations suggest sporadically structured firing of individual PNs at relatively short timescales under GluN1-Ab (**Fig. 1D** top). This was further corroborated by several findings (**Figs. 1I, S1D,E**), including the reduced correlation between the CV and median of ICIs (**Fig. 1I**) and the higher fraction of PNs having positive correlation between successive ICIs (**Fig. S1E**).

### GluN1-Ab enhances PN recruitment during synchronous events despite reduced network activity

Next, we investigated the influence of altered firing properties on network activity during SWS-like sleep. To this end, we analyzed Φ(t) as the fraction of active PNs over time (**Fig. 1D**, middle). The power spectrum of Φ(t) showed a distinct peak at ∼0.05–0.5 Hz in both groups (**Fig. 1J)**. However, the activity amplitude at this preferred oscillation frequency was higher under GluN1-Ab (**Fig. 1J**, green area), suggesting enhanced network rhythmicity. This was corroborated by the heightened variability of Φ(t) under GluN1-Ab (**Fig. 1K**), implying sporadic surges in network activity (**Fig. 1D**). In contrast, GluN1-Ab decreased the baseline network activity, evidenced by the lower Φ(t) power in the <0.05 frequency-band (**Fig. 1J**, purple area) and the Φ(t) distribution (**Fig. S1F**). We next assessed how GluN1-Ab affects synchronous events (SE), extracted as those activity periods in Φ(t) which significantly surpass detection threshold, determined separately for each FOV by randomizing CaT events per PN (Graf et al., 2022). We found a decrease in SE occurrence rate (**Fig. 1L**) without a change in the detection threshold or the duration of SEs (**Fig. S1G,H**). However, SE size (i.e. the fraction of PNs activated per SE) increased (**Fig. 1M**), and individual PNs were more frequently engaged in SEs under GluN1-Ab (**Fig. 1N**). Together, these results imply that GluN1-Ab reduces overall network activity while increasing the synchronization capacity of PNs, thereby facilitating hypersynchrony.

### Enhanced functional neuronal coupling under GluN1-Ab

We hypothesized that a strengthening of functional neuron-to-neuron connectivity (coupling) might underlie hypersynchrony under GluN1-Ab, despite overall reduced activity levels. Therefore, we calculated the pairwise correlation of binary CaT timeseries (**Fig. 1D**, top) using the spike-time tiling coefficient (STTC; **Fig. 2A-F**), as a frequency-independent measure of functional coupling (Cutts and Eglen, 2014). GluN1-Ab induced an overall increase in STTC, particularly in positively correlated PN pairs (**Fig. 2B**, **2D** top). This increase was more evident in pairs with significantly higher-than-chance correlation (**Figs. 2C,D** bottom and **S2A**), while the fraction of such pairs remained unchanged (**Fig. S2B**). Analyzing STTC relative to the inter-PN distance revealed increased coupling between both nearby PNs and those farther apart (**Fig. 2E**). We next assessed the dependency of this effect on the temporal range considered for measuring the correlation (tiling window). This analysis showed a robust increase in coupling of the significant pairs under GluN1-Ab over a range of timescales spanning more than an order of magnitude (**Figs. 2F, S2C**). Of note, shared input (e.g. onto a PN pair) may be a confounder when using STTC. Hence, to effectively control for the potential additive correlation, we utilized the partial correlation coefficient (PCC; **Fig. 2A**) (Meamardoost et al., 2021), where the coupling results (**Figs. 2G, S2D**) were qualitatively similar to those obtained using STTC. Together, our data suggest the enhanced functional coupling of a subset of CA1-PNs over extended spatial and temporal scales as a mechanism of hypersynchrony under GluN1-Ab.

**Figure 2.**
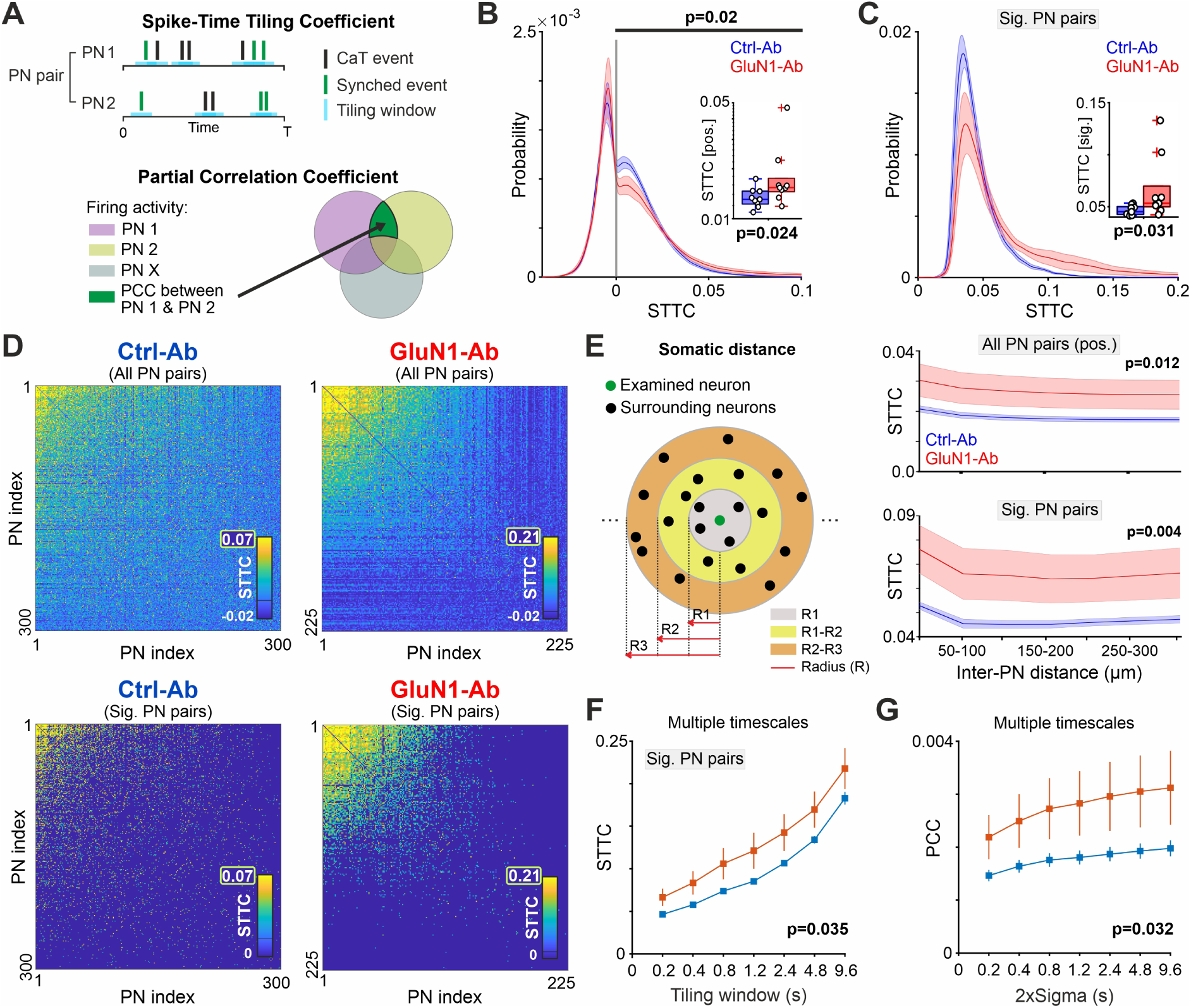
Enhanced functional neuronal coupling under GluN1-Ab. (**A**) Schematic representation of the STTC and PCC quantifications of coupling between firing-event trains of PN pairs. (**B**) Distribution of STTC indices, considering all PN pairs (n = 9 mice/4729 cells in Ctrl-Ab; n = 9 mice/3200 cells in GluN1-Ab; tiling window of 0.2 sec). For boxplots, only pairs with positive (pos.) correlation were considered. (**C**) Increased coupling of PN pairs with significantly high STTC (see Fig. S2A) under GluN1-Ab. Same format as in (B). (**D**) Example STTC matrices (re-ordered) of all (top row) and significantly (bottom row) coupled PN pairs from two individual FOVs. (**E**) Relationship between STTC and Euclidean somatic distance of PNs. Left, schematic of inter-PN distance used for quantification of the relationships shown in the right panels. For example, for 50-100 μm, for each PN, its average correlation with neurons locating within this range are computed. Radius: ca. 50 to 350 μm. (**F,G**) Increased neuronal coupling under GluN1-Ab at various timescales. The measures were quantified at multiple tiling windows of STTC (F) or standard deviation of Gaussian kernel used for smoothing CaT-trains in PCC (G). Curves presented as mean ± SEM. For statistical information, see Table S1.

### GluN1-Ab enhances functional PN clustering despite reduced global connectivity

To understand how this selective strengthened coupling (**Fig. 2**) can mediate the hypersynchrony over CA1, we hypothesized that GluN1-Ab may prompt an adaptive functional network reconfiguration which enhances neuronal clustering. Therefore, we resorted to the complex network analysis of (Rubinov and Sporns, 2010), in which each PN is considered as a node in a graph, and the link (connection) between each pair of nodes is determined by their coupling strength (STTC or PCC). We found that the node degree, defined as the number of links connected to a node, was decreased under GluN1-Ab (**Figs. 3A-C** and **S3A,B**), reflecting a lower number of functional connections between PNs. However, the right-tail of node degree distribution, including the dense nodes (i.e. highly connected), was preserved under GluN1-Ab (**Fig. S3A,B**). In addition, we found an increase in the clustering coefficient of PNs (i.e. the likelihood that the neighbors of a given node are also interconnected; **Fig. 3D**) as well as in different network-centrality measures (**Figs. 3SC**), implying enhanced functional PN-network clustering with strengthened hub-like properties of PNs under GluN1-Ab (**Fig. 3B,C**). Motivated by this finding, we next performed neuronal assembly detection analysis (Li et al., 2010; Graf et al., 2022) to assess whether enhanced clustering is attributed to a higher number or larger neuronal clusters. For this analysis, we utilized STTC instead of PCC, as it captures both direct and indirect coupling of neurons. GluN1-Ab increased the number of functional clusters, while having no effect on the total number of PNs assigned to the clusters relative to those without any cluster affiliation (**Fig. 3E-G, S3D**).

**Figure 3.**
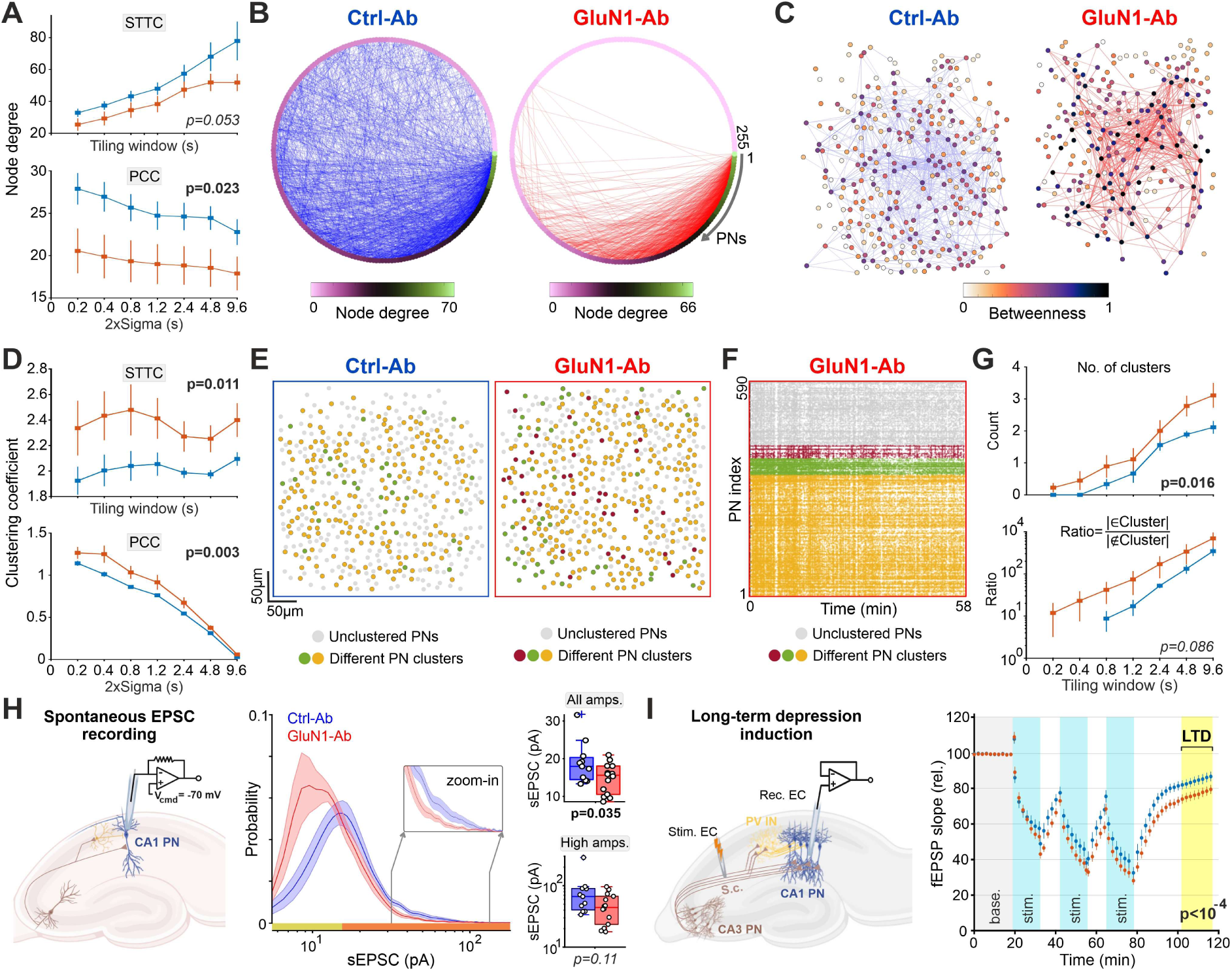
Enhanced functional neuronal clustering under GluN1-Ab. (**A**) Overall reduction in node degree under GluN1-Ab over different timescales, quantified based on STTC (top) and PCC (bottom) indices. Same format as in Fig. 2F,G. n=9 mice per group. (**B**) Circular graphs with links (connections) and nodes (PNs) of two example FOVs. Nodes were ordered in descending manner based on the ND of PNs, and links (unweighted) were drawn based on STTC matrices (tiling window of 0.3 s). To ease visualization, only links with a STTC above 95^th^ percentile (separately for each FOV) were shown. (**C**) Connectivity patterns of example graphs in (B). Each dot is a color-coded node (PN) based on its betweenness centrality (the extent to which a PN acts as a go-between for other PNs). To ease visualization of hub-like PNs (darker dots), the values were normalized to the maximum betweenness of the two FOVs. Line thickness encodes the STTC-based connection strengths, normalized to maximum STTC for each FOV separately. (**D**) Increased clustering coefficient of PNs under GluN1-Ab. (**E**) Example FOVs with detected PN clusters. (**F**) Rasterogram of the FOV shown in (E) under GluN1-Ab, with color-coded and re-arranged CaT-trains of PNs based on their affiliated cluster. (**G**) Number of detected PN clusters (top) and the ratio of PNs in clusters to unclustered PNs (bottom), per FOV. (**H**) Intracellular recording of sEPSCs from CA1-PNs in hippocampal slices. Left, schematic of recording configuration. Middle, distribution of sEPSC amplitude. Right, the median of all sEPSC amplitudes (top) and those above 95^th^ percentile (bottom), computed separately per neuron (n = 6 mice/11 cells in Ctrl-Ab; n = 6 mice/14 cells in GluN1-Ab). (**I**) Stronger long-term depression (LTD) under GluN1-Ab. Left, schematic of recording configuration. Right, slope of field EPSP relative to that of baseline period (gray). Yellow-colored period was analyzed for between-group difference after LTD induction (n = 6 mice/16 slices in Ctrl-Ab; n = 7 mice/21 slices in GluN1-Ab). Curves presented as mean ± SEM. For statistical information, see Table S1.

These results suggest that in response to generally reduced network activity under GluN1-Ab, the formation of functional PN sub-clusters is fostered in CA1. This occurs through an adaptive reconfiguration of functional connections, accompanied by a decrease rather than an increase in their overall count.

### Selective maintenance of effectively strong excitatory synapses under GluN1-Ab

How can the enhanced functional coupling and clustering be feasible while GluN1-Ab induces an overall decrease in the spontaneous excitatory postsynaptic currents (sEPSCs) onto CA1-PNs *ex vivo* (Ceanga et al., 2023)? We hypothesized that the maintenance of some sufficiently strong excitatory synapses between CA1-PNs provides the synaptic basis for this phenomenon. By re-analyzing our previously published sEPSCs data (Ceanga et al., 2023), we noted that despite an overall reduction in sEPSC amplitude under GluN1-Ab, the right tail of the distribution (comprising strong synapses) was largely preserved (**Fig. 3H**), hence supporting our hypothesis. Moreover, we wondered whether a disrupted LTD under GluN1-Ab is involved in the overall node degree reduction (**Fig. 3A**). While the reported impaired NMDAR-dependent LTP (Planagumà et al., 2016; Papi et al., 2024) may just elucidate the observed sparsity of strong sEPSCs (**Fig. 3H**, distribution right-tail), it does not solely suffice to explain the overall shift of sEPSC toward lower values. We therefore conducted an NMDAR-dependent protocol for LTD induction in Schaffer-Collateral synapses onto CA1-PNs in hippocampal slices *ex vivo* (Ahmed et al., 2011) (**Fig. 3I**, left). We found ∼20% reduction in synaptic strength under Ctrl-Ab (**Fig. 3I**). Importantly, this depression effect was stronger by ∼10% under GluN1-Ab (**Fig. 3I**), pointing to its role in the reduced number of the functional connections.

### Enhanced short timescale neuronal communication and spatiotemporal pattern similarity under GluN1-Ab

How do these altered functional and activity properties affect intrinsic neural coding in CA1 during SWS-like sleep? We quantified the similarity between binary spatial patterns of SEs based on their shared active PNs, using matching index (MI; **Fig. 4A**) and extracted significant MI values by comparing to randomized patterns (Romano et al., 2015; Graf et al., 2022). Despite reducing SE rate (**Fig. 1L**), GluN1-Ab skewed the distribution of significant MIs towards higher values, alongside a robust increase in MI across different thresholds, used to set the minimum fraction of active PNs per pattern (**Figs. 4B,C, S4A**). Secondly, to check the robustness of this finding, we detached this analysis from our SE detection method by repeating it for the binary spatial patterns obtained by dividing the entire recording time to non-overlapping bins. We again observed an increased overlap of patterns under GluN1-Ab, when using a bin-size of about ∼1 second (**Fig. 4D**), covering the range of SE durations (**Fig. S1H**). This effect was absent at longer bins (**Fig. S4B**), suggesting a distinct information processing role for these spatially-distributed neural patterns temporally confined to the timescale of SE durations.

**Figure 4.**
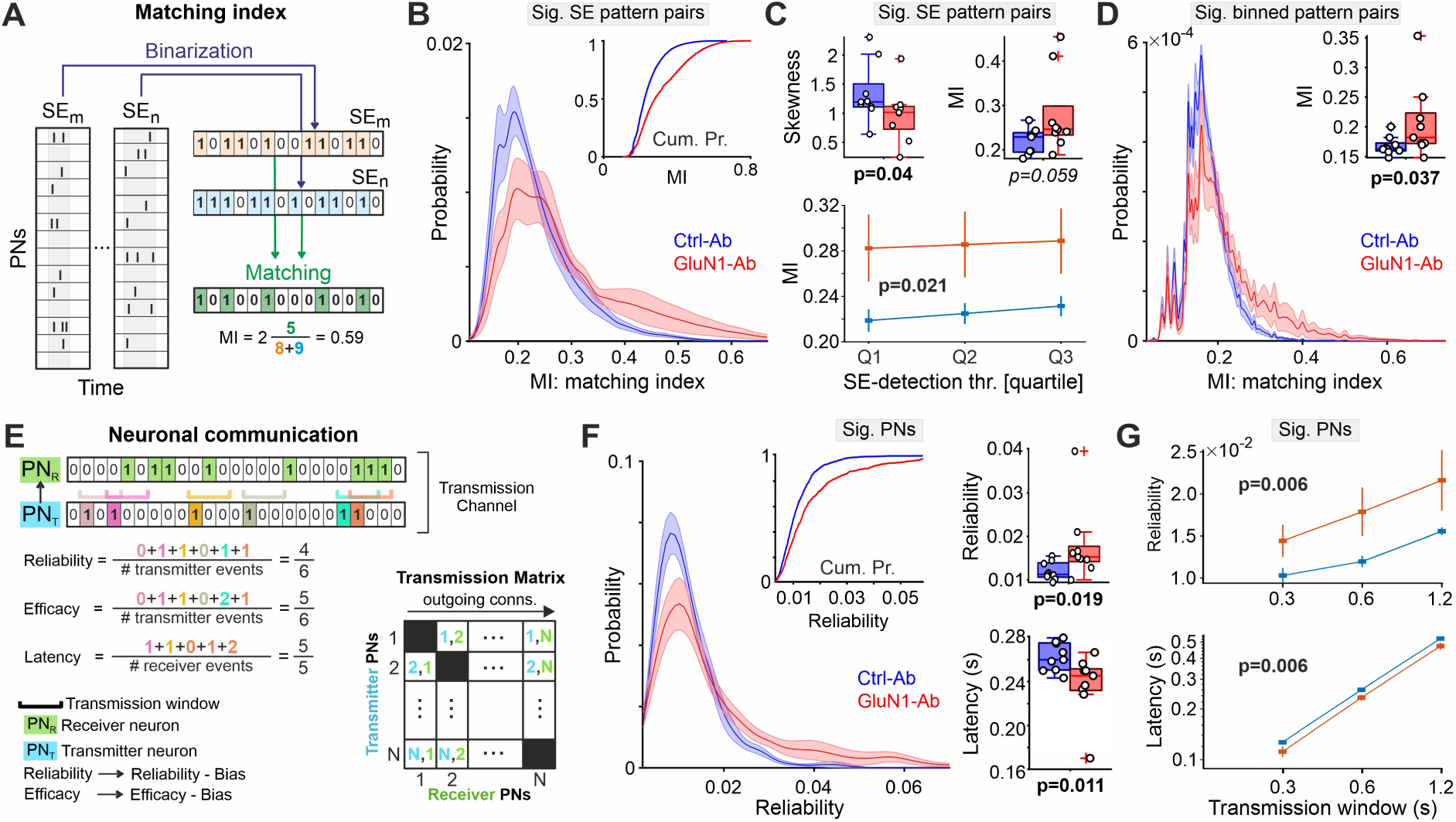
Enhanced short timescale neuronal communication and spatiotemporal patterns’ similarity under GluN1-Ab. (**A**) Schematic representation of similarity quantification between a SE pattern pair using MI measure. (**B,C**) Enhanced similarity between SE spatial pattern pairs with significantly high MIs, compared to similarity between spatially randomized SE patterns. n=9 mice per group. (B) Distribution of significant MIs. Same format as in Fig. 1E. (C) Top, boxplots of summary statistics of significant MIs. Bottom, significant MIs as a function of minimum number of active PNs in each SE pattern. Quartiles of the SE-detection thresholds of all mice: Q1≈3.9, Q2≈4.3, Q3≈4.9%. (**D**) Enhanced similarity between the spatial patterns, obtained by a non-overlap binning of entire recording time using a relatively short, fixed window size (∼1s). The binned pattern pairs with significantly high MIs were shown, similarly to (B). Inset: boxplot of corresponding median values. (**E**) Schematic representation of neuron-to-neuron communication parameters for a single PN_T_→PN_R_ transmission channel based on their CaT-onset trains. Bias denotes the randomization-based value of each parameter. For each PN, per parameter, its value is determined as the mean/median over its outgoing connections, i.e. over its corresponding row in depicted transmission matrix. (**F**) Enhanced reliability and reduced latency of individual PNs’ transmission to others. Per parameter, PNs with significant level (e.g. of reliability), as compared to randomized data, were considered (sig. PNs). Transmission window was set to 0.6 s. (**G**) Transmission parameters of sig. PNs at different plausible timescales. Curves presented as mean ± SEM. For statistical information, see Table S1.

Motivated by this finding, we investigated GluN1-Ab effect on neuronal communication throughout the entire recording period (**Fig. 4E**) (Ermentrout et al., 2008; Budak and Zochowski, 2019; Andrade-Talavera et al., 2023). We considered each PN as a transmitter (PN_T_) and any other PN as receiver (PN_T_→PN_R_), and quantified transmission reliability (fraction of successfully transmitted CaTs), latency (average time delay of this transmission), and efficacy (average number of CaTs triggered upon PN_T_ activation at PN_R_) within a determined transmission time-window (**Figs. 4E**), and extracted the PNs with a significant value as compared to randomized data (**Fig. S2A**). These PNs exhibited higher transmission reliability and efficacy with reduced latency under GluN1-Ab (**Figs. 4E-G, S4C-D**), while their fraction did not change (**Fig. S4E**). Together, these findings indicate selective enhancement of neuronal communication between PNs under GluN1-Ab.

## Discussion

What changes in the functional building-blocks at multi-neuronal level do mediate the NMDAR-Ab-induced network dysfunction? Here, using an established anti-NMDAR encephalitis mouse model, we provided new mechanistic insights by investigating how long-term exposure to GluN1-Ab alters internally generated hippocampal activity in CA1 during slow-wave sleep-like conditions minimizing the influence of sensory inputs. Our data show that GluN1-Ab renders PN-network generally hypoactive and concomitantly enhances neuronal coupling between a subset of CA1-PNs, accompanied by a profound functional network reconfiguration towards enhanced PN clustering. This reconfiguration together with the selective enhancement of neuron-to-neuron communication, in turn, promotes the spatial range of coordinated network activity (hypersynchrony) and similarity of spatiotemporal patterns at relatively short timescales. These indicate that anti-NMDAR encephalitis alters both the functional backbone of network configuration and intrinsic information transfer mechanism in hippocampal output microcircuits.

How can the network exhibit hypersynchrony, while baseline network activity is suppressed under GluN1-Ab? Firstly, this suppression possibly arises from the reduced NMDAR (Hughes et al., 2010; Planagumà et al., 2016; Ladépêche et al., 2018; Steinke et al., 2023) and AMPAR excitatory currents (Wright et al., 2021; Ceanga et al., 2023), inducing E-I-imbalance towards inhibition dominance. In particular, the possible CA1-PN hypoactivity aligns with recent *ex vivo* reports under GluN1-Ab, showing reduced excitatory synaptic input onto CA1-PNs; e.g. through CA3→CA1 (Ceanga et al., 2023). Secondly, corresponding theoretical models proposed that disinhibition of neural networks (Ceanga et al., 2023) or their exposure to instability and pathological dynamics (Rosch et al., 2018; Wright et al., 2021; Ceanga et al., 2023) underlie the induced network hypersynchrony. Beyond these modeling predictions, our empirical data unveiled a new mechanism: GluN1-Ab enhances the functional coupling and clustering among PNs thereby making the recruitment of additional neurons in coordinated network activities more likely, despite reduced baseline network activity. Indeed, this aberrant functional rewiring can render the neural networks more susceptible to epileptic seizures (Bragin et al., 2000; Wenzel et al., 2019) and disturb synaptic plasticity during non-REM activity which, in turn, may underlie severe learning and memory deficits of patients with anti-NMDAR encephalitis (Dalmau et al., 2017; Papi et al., 2024). Moreover, similar coexistence of low network activity and relatively large synchrony have been observed e.g. in several studies of Alzheimer’s disease (Busche and Konnerth, 2015; Klee et al., 2020), PCDH19-epilepsy syndrome (Giansante et al., 2023; Kowkabi et al., 2024), and in immature networks (Rahmati et al., 2017; Graf et al., 2022).

Our finding of the enhanced functional coupling may appear nontrivial given the disrupted LTP *in vitro* (Planagumà et al., 2016; Dalmau et al., 2017). However, the maintenance of a portion of effectively large sEPSCs, despite the overall reduction, supports the existence of a CA1-PNs subset with relatively strong synaptic coupling. This finding is corroborated by our functional topology analysis showing the augmentation of hub-like properties in a subset of PNs. The question arises of why these PNs did not follow the overall trend. Our finding of reduced latency of synaptic transmission as well as the decreased jitter of CA1-PN firing and integration time-window under GluN1-Ab (Ceanga et al., 2023) may hint at a shortened time-window of synaptic potentiation due to NMDAR hypofunction (Ceanga et al., 2023; Steinke et al., 2023; Papi et al., 2024), which in turn exposes synaptic connectivity of PNs with longer co-firing timescale to LTD while preserving the rest. This view is supported by our observed stronger LTD, and overall reduction in the number of functional connections except for a subset of PNs, under GluN1-Ab. Hence, a competitive impact of these altered plasticity mechanisms on the synaptic weights (Stanton, 1996; Diamond et al., 2005) might have caused the functional network reconfiguration and redistribution of sEPSC amplitudes. Moreover, these data may point to differential effect of GluN1-Ab on functional connectivity at local neuronal microcircuits and broader networks (Peer et al., 2017; Kuchling et al., 2024).

Importantly, such aberrant functional rewiring has been linked to the neuropsychiatric symptoms such as cognitive impairment, hallucinations, and perceptual distortions in schizophrenia (Whitfield-Gabrieli et al., 2009; Balu, 2016; Hamm et al., 2017; Silverstein and Lai, 2021). Furthermore, the higher similarity between the internally-generated spatiotemporal patterns observed in our data may contribute to the recently reported reduction in serial dependence (i.e. a readout of passive information maintenance across trials) in anti-NMDAR encephalitis patients (Stein et al., 2020), through entangling of these abnormal patterns and working memory traces. In addition, our observed CA1 network dysfunctions may contribute to the deficits in learning and memory consolidation (Paoletti et al., 2013; Dalmau et al., 2017; Peer et al., 2017; Papi et al., 2024), as our data relate to non-REM sleep-like state subserving these mechanisms (Wilson and McNaughton, 1994; Kim et al., 2019; Tamaki et al., 2020). This view is further reinforced as I) our data pertain to the CA1’s *stratum pyramidale*, which is crucial for hippocampal-cortical interactions involved in memory mechanisms (van Strien et al., 2009; Tang et al., 2017); II) we found higher number of internal, functional CA1-PN assemblies under GluN1-Ab which may be detrimental to ‘neural replays’ during memory consolidation (Wilson and McNaughton, 1994; Foster, 2017; Lewis et al., 2018).

Together, our findings reveal a profound functional rewiring of internal building-blocks of hippocampal computations which not only makes the network intrinsically susceptible to hypersynchrony but also alters its intrinsic information processing during slow-wave sleep-like conditions, thus providing new mechanistic insights into the NMDAR-hypofunction consequences and the pathomechanisms of anti-NMDAR encephalitis symptomatology.

## Acknowledgments

We thank Claudia Sommer for expert technical support. We thank Mehrtash Souri for computational assistance. This work was funded by the Deutsche Forschungsgemeinschaft (DFG research unit FOR3004; GE2519/8-1 and GE2519/9-1 [to C.G.], KI 1816/5-1 #432559020, KI 1816/9-1 #415914819 [to K.K.]), the Thüringer Aufbaubank (2017 FGI 0020 [to K.H.]), the German Federal Ministry of Education and Research (01GM1908B and 01EW1901 [to C.G.]), the Interdisziplinäres Zentrum für Klinische Forschung (IZKF) Jena (to M.C. and H.H.), the Foundation “Else Kröner-Fresenius-Stiftung” within the Else Kröner Research School for Physicians “AntiAge” (to M.C.), and the Schilling Foundation (to C.G.). Schematic figures of experiment setups in Figs. 1A, 3H-I were created using <u>BioRender</u>.

## Author contributions

V.R., M.C., K.K., K.H. and C.G. conceived and designed the study. V.R. analyzed data and prepared figures. V.R. and C.G. drafted the manuscript. M.C. conducted electrophysiological recordings. J.G. conducted two-photon imaging. V.R., M.C., J.G, K.K., and C.G performed data interpretation with inputs from all authors. V.R. and C.G. wrote the manuscript with inputs from all authors. C.G., K.K. and K.H. acquired funding. C.G. supervised this study.

## Supplementary Figures

**Figure S1.**
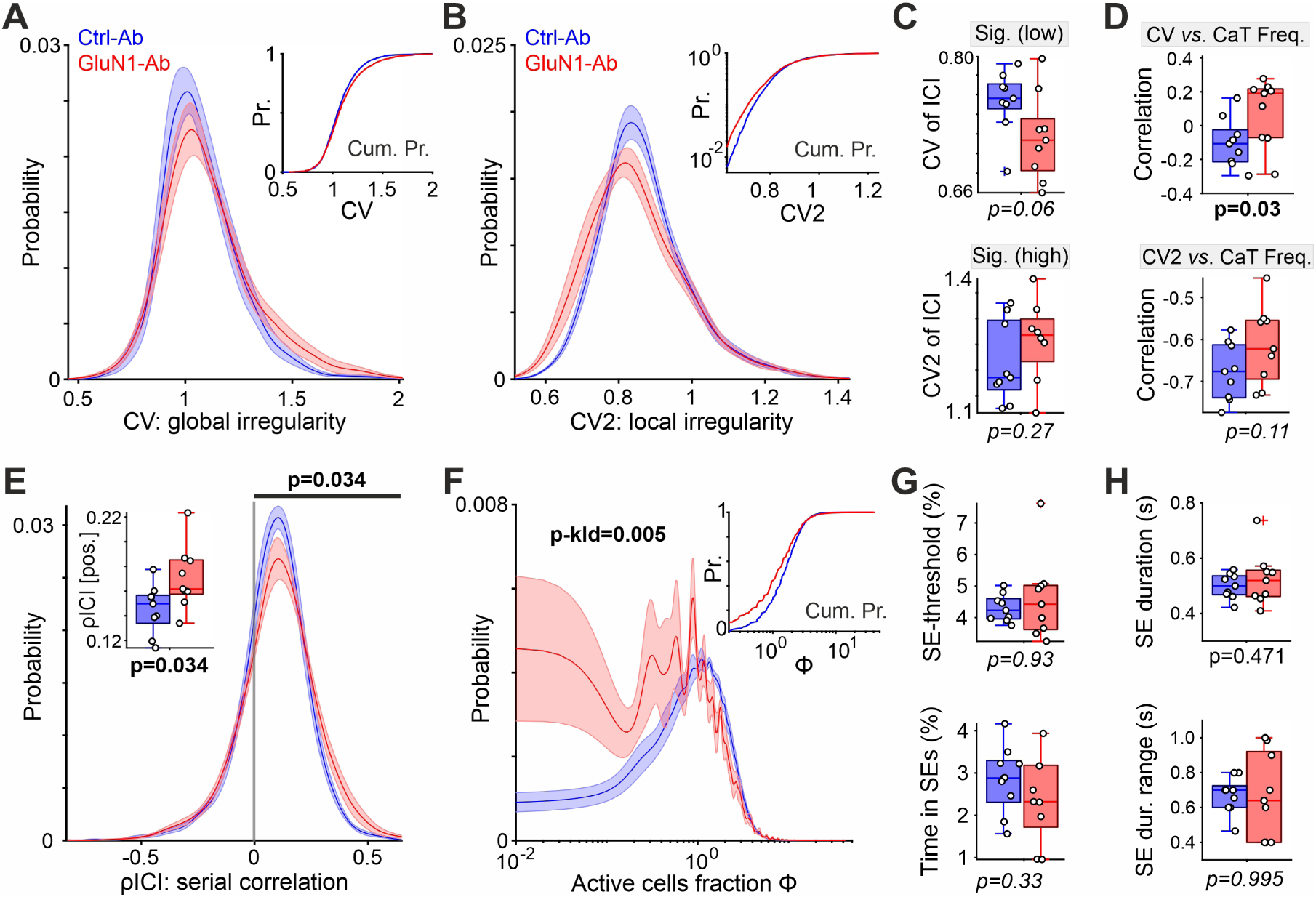
Complementary results related to Figure 1. (**A,B**) Distributions of global (A) and local irregularity of PNs’ ICIs (B). Same format as in (Fig. 1E). n = 9 mice/4729 cells in Ctrl-Ab; n = 9 mice/3200 cells in GluN1-Ab. (**C**) Unchanged irregularity level of PNs with significantly low CV and high CV2 values, as compared to randomized CaTs per PN. (**D**) The relationship between ICIs’ global irregularity (CV) and CaT frequency of PNs (top), and between ICIs’ local irregularity (CV2) and CaT frequency of PNs (bottom). (**E**) Redistribution of serial correlation of PNs’ ICIs (ρICI) towards higher values under GluN1-Ab, reflecting reduced PNs’ local firing irregularity. Inset: boxplots of mean of positive ρICI per mouse. (**F**) Redistribution of network activity towards lower values. Same format as in Fig. 1E. (**G**) SE-detection threshold (top) and the fraction of total duration of SEs relative to the entire recording time (bottom). (**H**) The mean duration of SEs (top) and the range of SE durations defined as the 95th *minus* 5th percentile of all SE durations per mouse (bottom). Curves presented as mean ± SEM. For statistical information, see Table S1.

**Figure S2.**
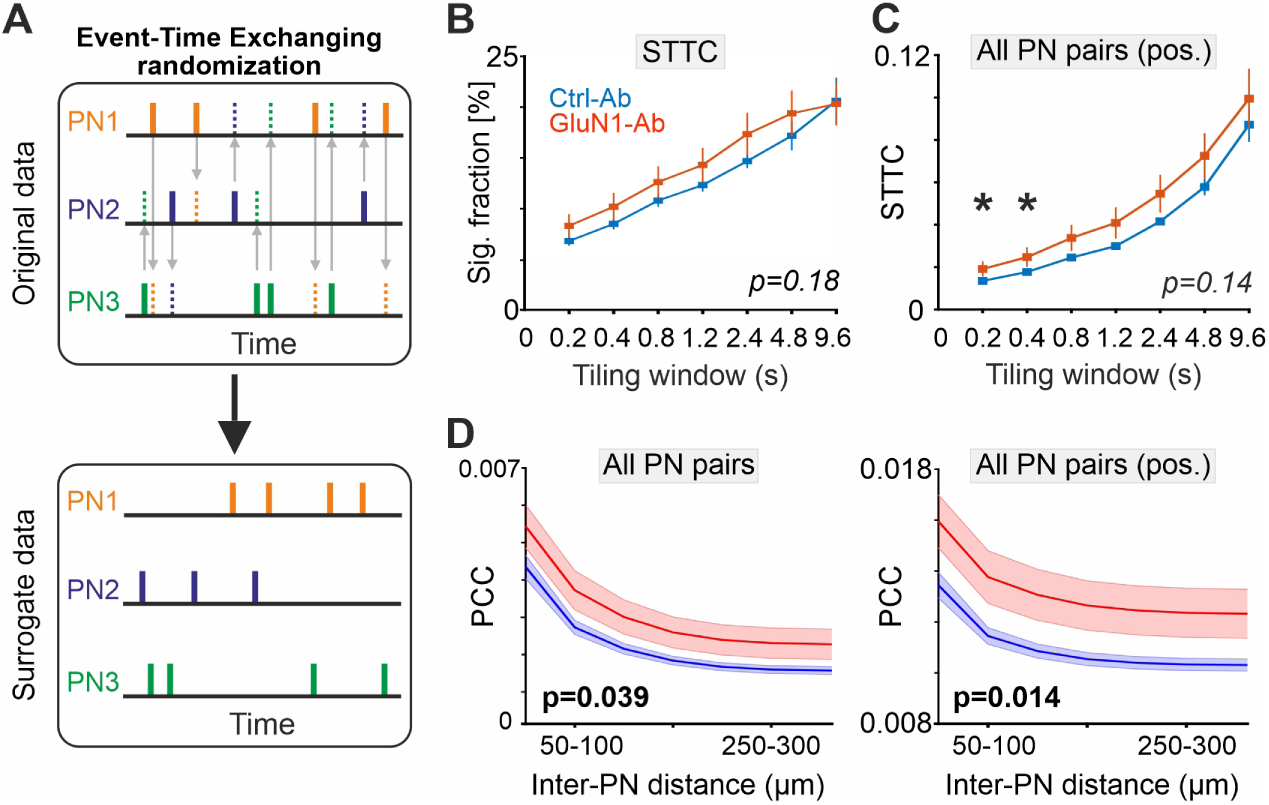
Complementary results related to Figure 2. (**A**) Schematic event-time exchanging (ETE) randomization to generate surrogate data to evaluate the significance level of the desired, empirical parameter value. In terms of pairwise correlation, the STTC between each of the PN pairs (here, three) is computed based on the empirical data (emp-STTCs). The ETE is performed and the STTCs are computed (sh-STTCs). This step is repeated M times, whereby a distribution of M sh-STTC values is built for each PN pair. The emp-STTC of each PN pair is compared to the extreme tail (e.g. 95^th^ percentile) of its corresponding sh-STTC distribution to evaluate its significance (see Methods for more detail). (**B**) Fraction of PN pairs with significantly high STTC, quantified at various timescales. Same format as in Fig. 2F. n = 9 mice per group. (**C**) Effect of GluN1-Ab on STTC of all PN pairs with positive value, at various timescales. Same format as in Fig. 2F. (**D**) Relationship between PCC and Euclidean somatic distance of PNs. Same format as in Fig. 2E. For statistical information, see Table S1.

**Figure S3.**
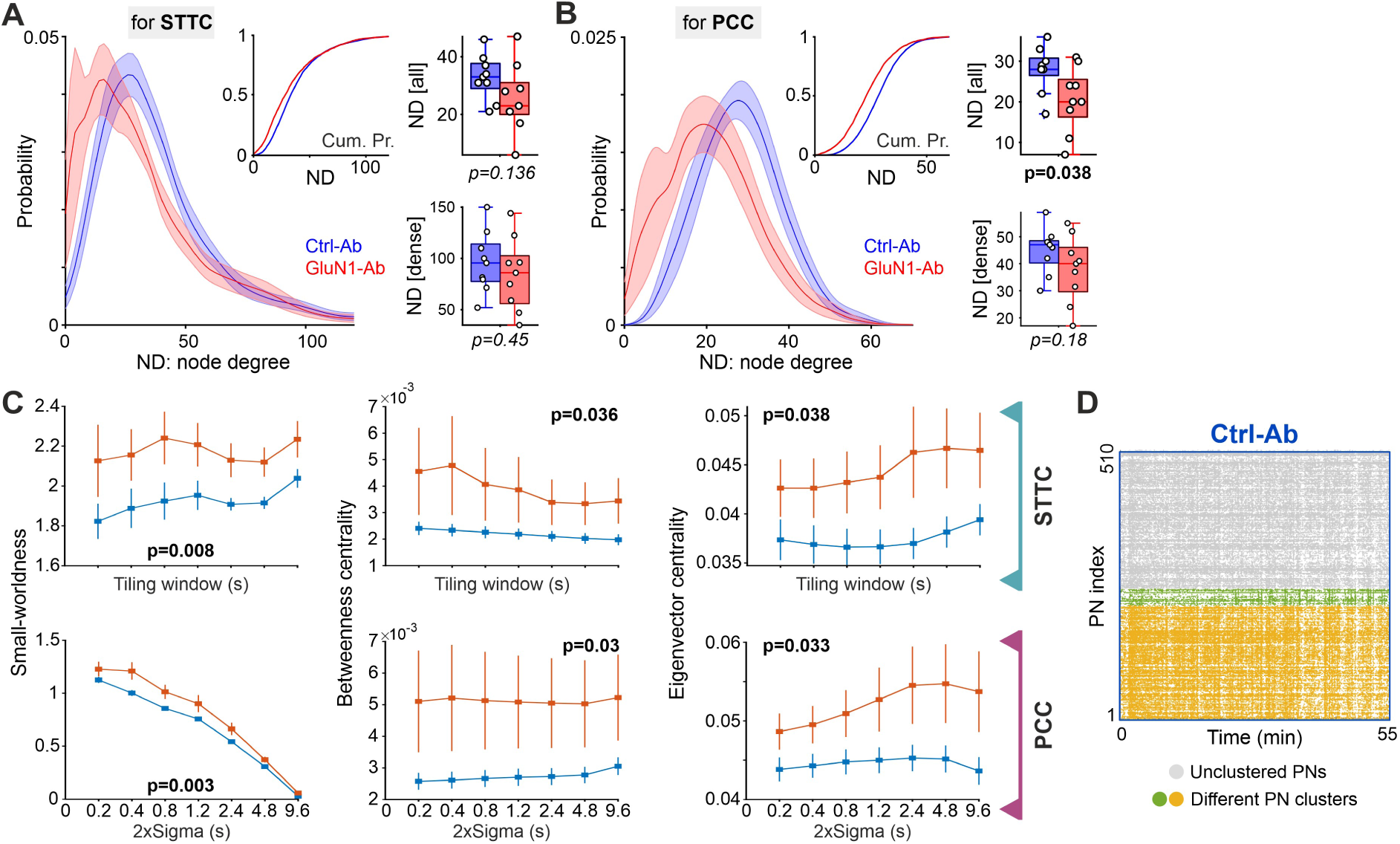
Complementary results related to Figure 3. (**A,B**) Overall reduction in the number of functional connections, despite the maintenance of a proportion of highly-connected PNs, under GluN1-Ab. Node degrees (NDs) of PNs were computed based on STTC (A) or PCC indices (B). Same format as in Fig. 1E. n = 9 mice/4729 cells in Ctrl-Ab; n = 9 mice/3200 cells in GluN1-Ab. Boxplots depict the mean ND of all PNs (top) and of those PNs with an ND above the 95^th^ percentile of distribution (dense nodes, bottom), computed separately per FOV. (**C**) Increased small-worldness (measuring the balance between local clustering and global connectivity in a network), betweenness centrality (measuring the extent to which a PN acts as a go-between for other PNs), and eigenvector centrality (measuring the importance of a node in a network based on its connections to other highly central nodes) under GluN1-Ab. Same format as in Fig. 2F,G. (**D**) Rasterogram of the FOV shown in Fig. 3E under Ctrl-Ab. Same format as in Fig. 3F. Curves presented as mean ± SEM. For statistical information, see Table S1.

**Figure S4.**
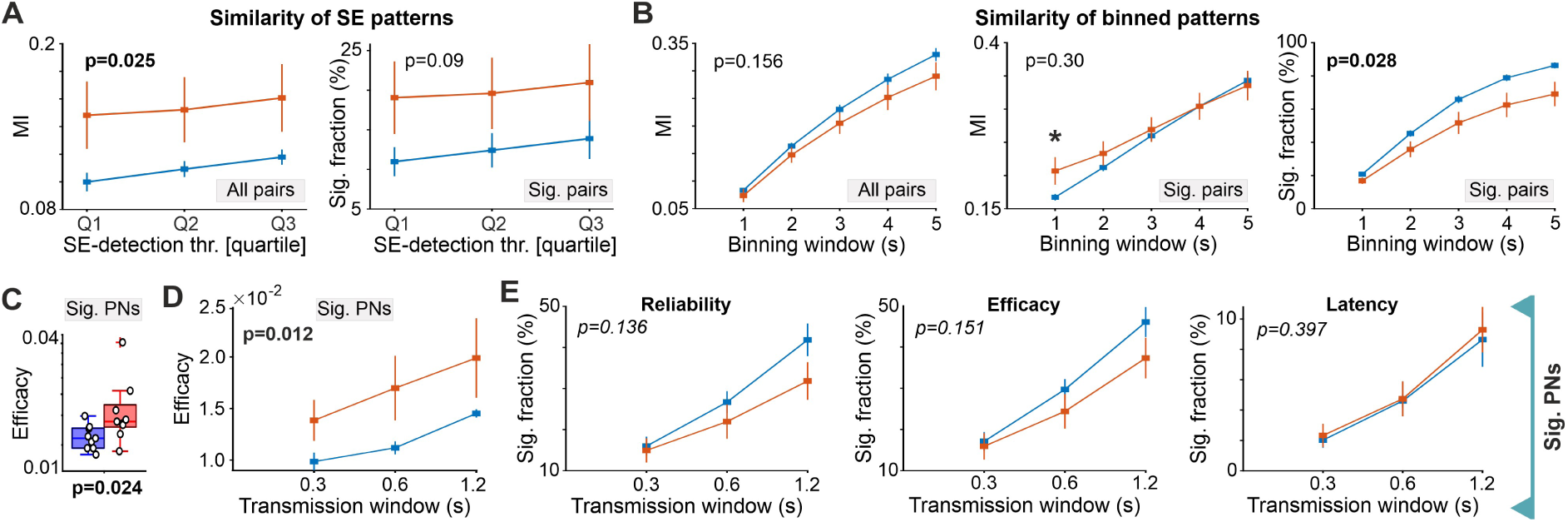
Complementary results related to Figure 4. (**A**) Left: enhanced similarity between SE binary patterns (all pairs) under GluN1-Ab. Right: Unchanged fraction of SE binary pattern pairs with significantly high MI under GluN1-Ab. Same format as in Fig. 4C bottom. (**B**) Similarity of spatial binary patterns as a function of window-size used for binning the recording time to extract the patterns. Left: for all pairs of binned patterns. Middle: for those pairs with significantly high MI. Right: fraction of these significant pairs. Note the elevated level of significant MIs at relatively shorter timescale (middle; see also 4D), and the decreasing trend in fraction of significant pairs at longer timescales (right). (**C**) Enhanced efficacy of individual PNs’ transmission to others (sig. PNs). Transmission window was set to 0.6 s (see also Fig. 4F). (**D**) Same as Fig. 4G, but for the transmission efficacy (sig. PNs). (**E**) Fraction of PNs with significant transmission reliability (left), efficacy (middle), and latency (right). Curves presented as mean ± SEM. For statistical information, see Table S1.

## Methods

### Resource availability

#### Lead contact

Further information and requests for resources and reagents should be directed to and will be fulfilled by the lead contact, Vahid Rahmati.

#### Materials availability

This study did not generate new reagents.

#### Data and code availability

Statistical data associated with this study is presented in **Table S1**. This paper does not report original code. MATLAB toolboxes for the Brain Connectivity Toolbox (<u>brain-connectivity-</u> <u>toolbox.net</u>), NoRMCorre (https://github.com/porteralab/NoRMCorre), UFARSA (https://github.com/VahidRahmati/UFARSA), FRACI (https://github.com/CABSEL/FARCI), FluoroSNNAP (https://www.seas.upenn.edu/~molneuro/software.html) as well as the MESc (https://femtonics.eu/) and Fiji (https://fiji.sc/) softwares are publicly available online. Any additional information required to reanalyze the data reported in this work paper is available from the lead contact upon request.

### Experimental model and study participant details

#### Animal model of NMDAR-encephalitis

Thirty three male C57BL/6J mice were housed under standard conditions at a controlled temperature (21 ± 1°C) and humidity (55 ± 10%) with 12-h illumination cycles, and food and water available *ad libitum*. Animal experiments were performed in accordance with the ARRIVE guidelines for reporting animal research and the experimental protocol was in accordance with European regulations (Directive 2010/63/EU) and was approved by the local animal welfare committee (Thüringer Landesamt für Lebensmittelsicherheit und Verbraucherschutz, 02–059/13 and UKJ-23-002), similarly to (Ceanga et al., 2023). Sixteen-weeks old (25-30g) mice were implanted with biventricular osmotic pumps (model 1002, Alzet, Cupertino, CA) with the following characteristics: volume 100 μL, flow rate 0.25 μL/h, and duration 14 days, as previously reported [25392198, 37768823]. The day before surgery the two pumps were each filled with 100μL of 1μg/μL human monoclonal NMDAR-Ab (#003– 102) or Control-Ab (#mGO53) (Kreye et al., 2016). Mice under isoflurane anesthesia were placed in a stereotactic frame, and a bilateral cannula (model 3280PD2.0/SP, PlasticsOne) was inserted into the ventricles (coordinates: 0.2 mm posterior and ±1.00 mm lateral from bregma, depth 2.2 mm). The cannulas were connected to two subcutaneously implanted osmotic pumps on the back of the mice.

### Method details

#### Surgical preparation for *in vivo* imaging

Thirty minutes before starting the preparation, 200 mg/kg metamizol (Novacen) was administered subcutaneously for analgesia. The animals were then placed on a warm platform and anesthetized with isoflurane (3.5% for induction, 1–2% for maintenance) in pure oxygen (flow rate: 1 l/min). A drop of eye ointment (Vitamycin) was applied to lubricate their eyes. The intraventricular cannula and osmotic pumps were carefully removed. For local analgesia, the skin was infiltrated with 2% lidocaine (s.c.). The scalp and periosteum were removed, and a custom-made plastic chamber with a central borehole (Ø 4 mm) was affixed to the skull using cyanoacrylate glue (UHU) (2.5 mm posterior from bregma and 2.2 mm lateral from midline). For the hippocampal window preparation (Mizrahi et al., 2004), the plastic chamber was securely attached to a preparation stage and superfused with warm artificial cerebrospinal fluid (aCSF) containing (in mM): 125 NaCl, 4 KCl, 25 NaHCO3, 1.25 NaH2PO4, 2 CaCl2, 1 MgCl2 and 10 glucose (pH 7.4, 35–36°C). A circular hole was drilled into the skull using a tissue punch (Ø 2.7 mm). The underlying cortical tissue and parts of corpus callosum were carefully removed by aspiration using a vacuum supply and a blunt 30G needle, taking care not to damage the alveus fibers. Once bleeding stopped, the animal was transferred to the microscope stage. During *in vivo* recordings, body temperature was continuously monitored and maintained close to physiological values (36–37°C) by means of a heating pad and a temperature sensor placed beneath the animal. Spontaneous respiration was monitored using a differential pressure amplifier (Spirometer Pod and PowerLab 4/35, ADInstruments). Shortly after transfer, Isoflurane was reduced to 0.6% (flow rate: 1 l/min). Recording of spontaneous activity commenced 60 min afterwards. At the end of each experiment, the animal was decapitated under deep isoflurane anesthesia.

#### Two-photon Ca^2+^ imaging *in vivo*

The recording chamber was continuously superfused with aCSF (as above). Stratum pyramidale, mainly comprising glutamatergic PNs, in the CA1 region was loaded with the membrane-permeable Ca^2+^ indicator Oregon Green 488 BAPTA-1 AM (OGB1, 500 µM, pipette resistance: ∼3 MΩ, pressure: 7 PSI, injection time: 1.5 min) using the multi-cell bolus-loading technique (Stosiek et al., 2003). We also positioned a tungsten microelectrode in the stratum radiatum of CA1, outside the recording field of view (FOV). After the aCSF was removed, the hippocampal window was filled with agar (1.5%, in 0.9 mM NaCl) and covered with a custom-made cover glass. Once the agar solidified, the chamber was reperfused with ACSF. To allow for de-esterification, recordings commenced approximately 60 minutes after OGB1 injection. Imaging was performed using an acousto-optic deflection (AOD) two-photon laser-scanning microscope controlled by the MES software (Femto3D ATLAS, Femtonics). Fluorescence excitation at 800 nm was provided by a tunable Ti:Sapphire laser (Chameleon Ultra II, Coherent) through a 20×/1.0 NA water immersion objective (XLUMPLFLN 20XW, Olympus). Emission light was separated from excitation light with a primary dichroic mirror (700 nm) and an IR blocker (700 nm) and finally detected by photomultiplier tubes (16 bit, H11706P-40, Hamamatsu). The high-speed arbitrary frame scanning mode of ATLAS enabled a fast frame rate of 38.81 Hz. The FOV was 329×329 µm at a pixel resolution of 0.9 µm/pixel (366×366 pixels, dwell time: ∼0.14 ns). Spontaneous activity in the dorsal CA1 was recorded for a total duration of 60 min. A single FOV was recorded per mouse. Data were acquired using MESc (Femtonics).

#### Analysis of imaging data

For each FOV, image stacks were registered using NoRMCorre (Pnevmatikakis and Giovannucci, 2017). For residual drift detection, a supporting metric was calculated as the Pearson correlation coefficient between the template image used for stack registration and the images of the registered image stack. Time periods with residual drift (e.g. abrupt mouse movement) were then visually identified (by inspecting the supporting metric and the registered image stack) and considered as missing values in subsequent analyses. We also removed 10 pixels from each end of FOV to avoid potential instability in alignment at the borders. For cell segmentation, we first smoothed the registered stack using a 3D Gaussian smoothing kernel (*imgaussfilt*3 Matlab function) of size [2 pixels, 2 pixels, 3 frames], followed by adapting the widely-used independent component analysis. The detected raw ROIs were then visually double-checked with the registered image stack, and the final ROIs were considered as the putative CA1-PNs’ somata. We then down-sampled the (non-smoothed) registered data to 9.7 Hz (∼100 ms) (Graf et al., 2022) to improve the signal-to-noise ratio of traces. For each ROI, we obtained the mean F(t) by frame-wise averaging across all pixels of that ROI in registered stack. The somatic fluorescence traces were obtained as relative changes from resting fluorescence levels (ΔF/F_0_). The resting fluorescence F_0_(t) was defined as the moving median over ∼30 seconds. For overlapping ROIs, we extracted the fluorescence trace of each ROI based on its non-overlapping pixels. For each ROI, CaT onsets were extracted from its ΔF/F_0_ trace using UFARSA, a general-purpose event detection routine (Rahmati et al., 2018). Reconstructed CaT onsets were translated into a binary activity vector (1 – event, 0 – no event) and used for the following analyses. In total, 10 FOVs were recorded for each group (1 FOV per mouse), where we had to exclude one FOV from each group due to the instability of recordings.

#### Long-term synaptic depression induction *ex vivo*

The protocol for inducing NMDAR-dependent LTD in hippocampal slices from adult mice was adapted from (Ahmed et al., 2011). Mice (n=6 for Ctrl-Ab, and n=7 for GluN1-Ab) were deeply anesthetized using isoflurane and the brain was quickly removed into ice-cold protective artificial cerebrospinal fluid (paCSF, in mM: 95 N-Methyl-D-Glucamine, 30 NaHCO3, 2.5 KCl, 1.25 NaH2PO4, 10 MgSO4, 0.5 CaCl2, 20 HEPES, 25 glucose, 2 thiourea, 5 Na-ascorbate, 3 Na-pyruvate, 12N-acetylcysteine, adjusted to pH7.3 and 300–310mOsmol, saturated with Carbogen). Transverse, 400µm-thick hippocampal slices (n = 6 mice/16 slices for Ctrl-Ab; n = 7 mice/21 slices for GluN1-Ab) were prepared in ice-cold paCSF on a vibratome (VT 1200S, Leica, Wetzlar, Germany) and transferred for warm recovery in paCSF at 33°C for 12 min. Thereafter, slices were transferred for recovery (at least 1h) at room temperature in aCSF^+^ (containing in mM: 125 NaCl, 25 NaHCO3, 2.5 KCl, 1.25 NaH2PO4, 1 MgCl2, 2 CaCl2, 25 glucose, 2 thiourea, 5 Na-ascorbate, 3 Na-pyruvate, 12 N-acetylcysteine, adjusted to pH 7.3 and an osmolarity of 300–310 mOsmol, and saturated with Carbogen). For recordings, slices were transferred in a custom-made interface-like recording chamber under continuous perfusion (3 mL/min, Ismatec, Wertheim, Germany) with 30°C aCSF containing in mM 125 NaCl, 4.5 KCl, 25 NaHCO3, 1.25 NaH2PO4, 2 MgSO4, 3 CaCl2, 10 glucose, saturated with Carbogen.

We were blinded to the experimental treatment of the animals during recording and data analysis. CA1-PNs were visually identified using a microscope (Examine.Z1, Zeiss, Jena, Germany) equipped with differential interference contrast optics. Recording and stimulation pipettes were pulled using a P-87 horizontal pipette puller (Sutter Instruments, Novato, CA, USA) from thick-walled borosilicate glass (0.86x1.50, Science Products, Kamenz, Germany) and were filled with aCSF. The field excitatory postsynaptic potential (fEPSP) was recorded and digitized with a MultiClamp 700B amplifier (Molecular Devices, Sunnyvale, CA, USA), and an Axon Digidata 1550B digitizer (Molecular Devices, Sunnyvale, CA, USA), respectively. Signals were low-pass filtered at 2kHz and digitized at 20kHz.

Stimulation of the Schaffer collaterals (SC) was performed with an aCSF-filled micropipette acting as a monopolar stimulation electrode that was placed 300µm away from the recording electrode. Stimulation was applied through a constant current stimulation unit (DS3, zigitimer). The LTD Stimulation protocol was conducted using the following steps. First, an input/output curve of field excitatory postsynaptic potential (fEPSP) responses to incremental SC stimulation was measured in the CA1 stratum radiatum by increasing stimulation strength from 20µA to 150µA delivered at 0.1Hz. Slices which did not generate a fEPSP amplitude of at least 1mV were discarded. Stimulation strength was adjusted for ∼50% (40-60%) of maximal fEPSC slope (mV/ms). LTD was induced after a baseline period of 20 minutes (stimulation at 0.05Hz) by three rounds of low-frequency stimulation (LFS) of 1500 pulses at 2Hz, with an interval of 10 minutes between each round (Ahmed et al., 2011). This protocol results in an NMDAR-dependent (non-selective to NMDAR-subunit) LTD that persists in aged mice. After LTD inductions fEPSP slope was recorded for a further 40 minutes at 0.1Hz. Analysis of recorded signals was performed in Clampfit 10.5 (Molecular Devices, Sunnyvale, CA, USA).

#### Properties of pyramidal neuron activity

For each PN, we quantified the temporally local irregularity level of its CaT onsets using CV2, as a local and relatively rate-independent measure of spike-time irregularity (Holt et al., 1996): 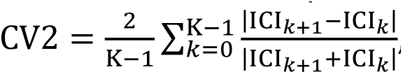, where ICI_k_ and ICI_k+1_ are the *k*th and (*k+1*)th inter-CaT intervals (ICIs) of the cell, and K is the total number of its ICIs. The global irregularity of each PN was computed using the coefficient of variation as CV = σ_ICI_⁄μ_ICI_, where σ_ICI_ and μ_ICI_ are the standard deviation and mean of the PN’s ICIs. To determine the PNs with a significant value of global irregularity, we compared the CV of each cell to those obtained based on its randomly shuffled CaT onsets (uniform distribution, 500 times). This randomization kept the mean CaT frequency of each cell unchanged. For each PN, we then computed the 1st and 99th percentile values of its surrogate data (500 values). This was followed by computing the median of these values over all PNs, separately for 1st and 99th percentiles, whereby determining the grand left-tail (η_L_) and right-tail (η_R_) significance levels of CV for all PNs. This approach enables setting a low and a high significance level of CV for the whole FOV. Finally, for each PN, we considered its empirical CV to be significantly low (resp. high) if its smaller (resp. bigger) than empirical η_L_ (resp. η_R_). Using similar approach, we extracted the PNs with a significantly low or high CV2 level. Additionally, for each PN, we estimated its ICI correlation (ρICI), as the correlation of consecutive ICIs using Spearman’s rank-order correlation of order one (Farkhooi et al., 2009). To achieve more robust results, PNs with less than ten ICIs were excluded for these measures.

#### Functional pairwise correlations

To compute the pairwise correlation between neural activity of PNs (functional connectivity) we used two measures: 1) spike-time tiling coefficient (STTC), and partial correlation coefficient (PCC). STTCs were computed for all possible PN pairs with a synchronicity window Δt using custom written Matlab code (Cutts and Eglen, 2014). STTCs derived from measured data were compared to those from event-time exchanging (ETE) surrogate data, generated by randomly exchanging the CaT onsets across PNs, thereby preserving both the mean CaT frequency of each PN and the sum network activity per time-point. This randomization was performed 500 times. For each pair, using its surrogates, we determined the significance of its empirical STTC (95th percentile). For synchronicity window, we set 2xΔt = 2, 4, 8, 12, 24, 48, or 96 frames, where 1 frame is ∼100 ms, and the tiling window is determined as [-Δt, Δt]. Note that STTC between a pair of PNs captures not only their direct coupling but also any indirect coupling induced by, for example, a third PN that sends input to both. Hence, to effectively control for this additive correlation whereby achieving a relatively unmixed pairwise correlation, we computed PCC by adapting FARCI toolbox (Meamardoost et al., 2021). The partial correlation PCC_ij_ between the *i*th and *j*th PNs was computed as 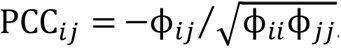 where the precision matrix ϕ = Σ^-l^, and Σ is NxN covariance matrix of neuronal activity among N neurons calculated based on the smoothed CaT-onset trains. For smoothing, we convolved each train with a Gaussian kernel with a standard deviation (Sigma) of 2, 4, 8, 12, 24, 48, or 96 frames. Both STTC and PCC matrices are symmetric and NxN in size, where their values range from -1 and +1 which indicate perfect negative and positive correlations, respectively.

#### Functional neuronal assembly

To detect neuronal assemblies (PN clusters) we subjected the pairwise correlation matrices (see above) to the eigendecomposition clustering method, by adapting FluoroSNNAP toolbox (Patel et al., 2015). Briefly, this method decomposes a given similarity matrix (here, STTC matrix) into a set of eigenvalues and eigenvectors. The number of significantly large eigenvalues determines the number of neuronal clusters, and their corresponding eigenvectors contain the information about cluster structure (i.e. the set of neurons belonging to each cluster). We used the same ETE surrogate data for testing the statistical significance (α = 5%) of the eigenvalues. This procedure enabled us to identify the clusters of PNs, which occurred beyond chance level. For more details about the clustering method and its mathematical description see (Li et al., 2010).

#### Functional network topology

To determine the properties of functional network topology (connectivity motifs) we applied the complex network analysis (Rubinov and Sporns, 2010) to thresholded STTC and PCC matrices computed above. To this end, for each FOV, we first binarized its empirical symmetric STTC (or PCC) matrix using the 95th percentile value of its ETE surrogate data. This was followed by subjecting the resulted binary (unweighted) and undirected matrix as the input to the functions implemented in the MATLAB Brain Connectivity Toolbox (Rubinov and Sporns, 2010), where each PN was considered as a node and the link (connection) between each pair of nodes was determined by their functional coupling (STTC or PCC). We quantified five topology metrics mainly relating to the clustering and centrality features of the network: I) node degree (ND), measuring the number of connections each PN has with other PNs, II) clustering coefficient (CC), measuring the likelihood that the neighbors of a given PN are also interconnected, III) eigenvector centrality (EVC), measuring the importance of a node in a network based on its connections to other highly central nodes, IV) betweenness centrality (BWC), measuring the extent to which a PN acts as a go-between for other PNs, and V) small-worldness (SW), measuring the balance between local clustering and global connectivity in the network. To this end, we quantified I)-III) using *degrees_und*, *clustering_coef_bu*, and *eigenvector_centrality_und* functions, respectively. For BWC, we applied *betweenness_bin* function to the output of *weight_conversion* function implementing ‘lengths’ conversion method. For SW, we first obtained characteristic path length (CPL) by computing the distance matrix (*distance_bin* function) and using it as the input to *charpath* function without including distances on the main diagonal and infinite distances. CC and CPL values were normalized by dividing them by corresponding “null” values, which were determined by generating 100 synthetic random networks (*makerandCIJ_und* function), computing the same parameters at each iteration, and then averaging them per parameter. To account for the different number of nodes (N) across networks, we divided the BWCs by [(N-1)*(N-2)]. Finally, we computed SW by dividing the normalized CC by the normalized CPL.

#### Network activity properties

For each FOV, we first computed network activity timeseries. To this end, in the CaT train of each PN, we set ±ωt frames around each CaT to 1 whereby accounting for some temporal jitter in the detection of CaT-onsets. Unless otherwise stated, ωt was set to 1. This was followed by computing the mean across the resulting CaT vectors of all individual PNs to obtain the empirical fraction of active cells per frame Φ(t) (Graf et al., 2022). To quantify the variability (fluctuations) in network activity we calculated the coefficient of variation of Φ(t). To assess the rhythmicity of network activity, we calculated the power spectral density (PSD) of Φ(t) using Welch’s method (MATLAB *pwelch()* function) with 20 s Hamming windows, 75% overlap. The PSD of each FOV was normalized to its total PSD to obtain a more robust measure of oscillatory bands when averaging over mice per group. To detect synchronous events (SEs) as the significant network co-activation periods, we randomly shuffled CaT onsets of all PNs (uniform distribution; 1000 times), computed the surrogate Φ(t) (as above), and defined the 99.99th percentile of all surrogate Φ(t) as the threshold for NB detection. The NB threshold was determined separately for each FOV, to account for different mean CaT frequencies. We then considered any frame with an empirical Φ(t) exceeding the threshold as belonging to an SE. In the resulting binary SE vectors, 0→1 transitions were defined as SE onsets and 1→0 transitions as SE offsets. Using the binary SE vectors, we extracted following parameters: (1) the SE occurrence rate as the number of SEs divided by the total available recording time, (2) the relative time the network spent in SEs, (3) the average SE duration (offset *minus* onset *plus* 1frame), (4) SE size as the fraction of PNs which were active in at least one frame of a given SE. In addition, we computed the synchronization capacity of each PN as the fraction of SEs in which it participated, divided by its CaT frequency to control for different PN activity levels.

#### Similarity of spatiotemporal patterns

To investigate the similarity of SE spatiotemporal patterns, we represented each SE period as a binary spatial pattern (vector) of active and inactive PNs of size Nx1, where N is the number of PNs in FOV. We then quantified the similarity between each two patterns Pat_i_ and Pat_j_ using the so-called matching index (Romano et al., 2015; Graf et al., 2022): 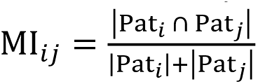. MI ranges from 0 (no similarity) to 1 (perfect similarity) and approximates the number of shared active PNs (i.e. common ones) between pattern pairs. For each pattern pair, we determined the significance of its empirical MI (99th percentile) using the surrogate data obtained by randomly shuffling the values within binary patterns (500 shuffles). In addition, to assess the patterns similarity as a function of SE size, we thresholded SEs based on their size, and recomputed the measure between the remaining SEs. For this, we used three thresholds obtained by calculating the quartiles of SE-detection thresholds (see above) of all FOVs: Q1≈3.9, Q2≈4.3, Q3≈4.9%. This is equivalent to using the same thresholds for all mice. To assess whether our findings in this analysis is specific to SE patterns or extends to the broader network activity, we divided the entire recording time to non-overlapping bins, converted them to binary vectors, and computed their similarity using the same procedure (250 shuffles, 99th percentile). For the binning, we used windows of size 10, 20, 30, 40, or 50 frames. For more robust results, in addition to silent bins, the patterns including less than five active PNs were also excluded.

#### Neuronal communication properties

To investigate the effect of GluN1-Ab on neuronal communication between neurons (Ermentrout et al., 2008; Budak and Zochowski, 2019; Andrade-Talavera et al., 2023), we analyzed the corresponding signal transmission properties based on the reconstructed binary CaT-onset trains of PNs. Per communication channel PN_T_→PN_R_ (T: transmitter, R: receiver; for brevity, we here refer to each CaT of PN_T_ as tCAT and of PN_R_ as rCaT), we approximated its transmission reliability, efficacy, and latency within a specified transmission time window (W_TR_) after each tCaT. The reliability quantifies the proportion of tCaTs that successfully transmitted to PN_R_ within W_TR_. It is defined as the ratio of tCaTs that have at least one corresponding rCaT to the total number of tCaTs. To avoid double (or multiple) counting, each rCaT is only considered for the closest tCaT within W_TR_. The reliability ranges from 0 to 1, where 1 indicates 100% success rate for transmission (of note, the corresponding transmission failure can be determined as: 1 *minus* Reliability). The efficacy is computed as the total number of rCaTs occurring within the W_TR_ after each tCaT divided by the total number of tCaTs. Each rCaT is only counted once, even if it falls within the window of multiple tCaTs, to avoid double counting. The efficacy ranges from 0 to potentially greater than 1, depending on the specific neural activity patterns. The Latency refers to the average time delay between a tCaT and the corresponding rCaT within W_TR_. The average latency is computed across all successful transmissions, with each rCaT only being counted once, by considering its latency to the closest tCaT. For the transmission window, we set W_TR_ = 3, 6, or 12 frames (1frame ≍ 100 ms), consistent with the timescale of SE durations in our data and of plausible short-term synaptic plasticity (Rahmati et al., 2017). For each FOV, this analysis provided us by a generally asymmetric NxN transmission matrix per parameter, where N is the number of PNs in FOV; note that the transmission properties of PN_x_→PN_y_ channel is generally different from PN_y_→PN_x_. We defined the reliability of each PN (as transmitter) by computing the median over its corresponding row, i.e. over its outgoing channels to all other PNs (as receivers), in the corresponding matrix. The efficacy and latency of each PN were quantified similarly. We considered the PNs with the empirical reliability and efficacy levels bigger than 95th, and the empirical latencies smaller than 5th, of their corresponding ETE surrogate data as significant, separately. When reporting the reliability and efficacy of each PN, we subtracted the mean of its corresponding surrogate values to account for the potential differences in PN CaT frequencies.

### Quantification and statistical analysis

Statistical analyses were performed using OriginPro 2019, MATLAB 2020a. All data are reported as mean ± standard error of the mean (SEM), if not stated otherwise. The Shapiro– Wilk test was used to test for normality. Parametric testing procedures were applied for normally distributed data; otherwise, nonparametric tests were used. The *F*-test was used to test for homogeneity of variances. Except for the Shapiro–Wilk test and F-test where p values <0.05 were considered statistically significant, the actual p values were stated for other tests. The skewness and Spearman’s rank correlation coefficient were computed using *skewness* and *corr* Matlab functions. Curve permutation tests were performed by comparing the area under the curve (AUC) of the difference of the group-mean (or median) curves, with those AUC values obtained after shuffling the individual curves across groups, with 1 million times repetition; see Ref. (Ceanga et al., 2023) for more details. The distance of group-mean (or median) probability distribution of GluN1-Ab group relative to the equivalent one of Ctrl-Ab (as reference) was measured using the well-known Kullback–Leibler divergence (KLD) metric. Similarly to the curve permutation, the p-value of the empirical KLD (p-kld) was computed by comparing to KLD values obtained after shuffling the individual curves across groups, with 1 million times repetition. This KLD permutation test was performed on the data with no significant change in the mean or median, to test the difference in their distribution shapes. Only for better visualization, the depicted probability distributions were smoothed using a normal kernel density estimation (*ksdensity* Matlab functions). Details of the applied statistical tests with the sample sizes are provided in **Table S1**.

